# Exometabolite dynamics over stationary phase reveal strain-specific responses to nutrient limitation

**DOI:** 10.1101/2020.06.05.137489

**Authors:** John L. Chodkowski, Ashley Shade

## Abstract

Microbial exponential growth is expected to occur infrequently outside of the laboratory, in the environment. Instead, resource-limited conditions impose non-growth states for microbes. However, non-growth states are uncharacterized for the majority of environmental bacteria, especially in regard to exometabolite production. To investigate exometabolite production in response to nutrient limitation, we compared exometabolites produced over time in stationary phase across three environmental bacteria: *Burkholderia thailandensis* E264 (ATCC 700388), *Chromobacterium violaceum* ATCC 31532, and *Pseudomonas syringae* pathovar *tomato* DC3000 (ATCC BAA-871). We grew each strain in monoculture and investigated exometabolite dynamics over time from mid-exponential to stationary phase. We focused on exometabolites that were released into the media and accumulated over 45 hours, including approximately 20 hours of stationary phase. In concert, we analyzed transcripts (RNAseq) to inform interpretation of exometabolite output. We found that a majority of exometabolites released under these conditions were strain-specific. A subset of identified exometabolites were involved in both central and secondary metabolism. Transcript analysis supported that exometabolites were released from intact cells, as various transporters were either upregulated or consistently expressed. Interestingly, we found that all strains released succinate, and that each strain re-routed their metabolic pathways involved in succinate production during stationary phase. Overall, these results show that non-growth states can also be metabolically active and dynamic. Furthermore, they show that environmental bacteria have the capability to transform a resource-limited extracellular environment into a rich chemical milieu. This work has implications for understanding microbial community interactions via exometabolites, and within resource-limited environments.

**Importance:** Non-growth states are common for bacteria that live in resource-limited environments, and yet these states remain largely uncharacterized in cellular metabolism and metabolite output. Here, we investigated and compared stationary phase exometabolites and RNA transcripts for each of three environmental bacterial strains. We observed that diverse exometabolites were produced and that they collectively exhibited clear and directional dynamics over time. Additionally, each bacteria strain had a characteristic exometabolite profile and dynamic. This work affirms that stationary phase is not at all “stationary” for these bacteria, and sets the stage for understanding how individual metabolisms support interspecies interactions in resource-limited environments.

## Introduction

Much of microbiology research in the laboratory is conducted with bacterial or archaeal cells that are growing exponentially. However, it is estimated that 60% of microbial biomass in the environment is in a non-growing state (1, 2). Various abiotic and biotic stressors are known to induce non-growth states, but perhaps the most common is resource limitation. Labile resources can be low in an environment, as characteristic of the oligotrophic open ocean, or they can be available but inaccessible, as typical in heterogeneous soil matrices. Thus, unlike most cultivated laboratory strains, environmental microbes experience short periods of accessible resources punctuated by long periods of famine (3, 4).

While Gram-positive bacteria can survive resource limitation through sporulation (5), Gram-negative bacteria can persist in stationary phase without entry into a specialized dormant cell structure (6). Instead, Gram-negative bacteria survive in stationary phase by employing various stress response adaptations (7). Stress response adaptations include changes to cell morphology, transcription, translation, and metabolism. Furthermore, in stationary phase, microbes can re-route metabolic pathways to maintain essential components of the cell and the proton motive force (8). While these adaptations are thought to serve as survival mechanisms, the levels and types of metabolic activities in stationary phase are not well understood for most environmental microbes.

It is known, however, that microbes can exhibit appreciable metabolic activity in stationary phase (9). For example, entry into stationary phase resulted in prolonged protein production in *Escherichia coli* despite that overall protein levels decreased (10). Metabolomic studies of *E. coli* in stationary phase support that there is continued metabolite production and transformations despite growth arrest (11–13). These studies have provided valuable insights into stationary phase physiology for model microorganisms. However, metabolome studies of model microorganisms have generally focused on the dynamics of intracellular metabolites. It is expected that understanding exometabolite dynamics can provide insights into metabolic responses that are relevant for microbial communities and interactions among community members that coexist in the environment.

Exometabolomics is the characterization of small, extracellular molecules either released or transformed by a microbe (14). Characterizing exometabolites can provide insights into the potential for microbes to engage locally with other microbes and the environment via release of small molecules (15). The effect of these small molecules on neighboring microbes can range from cooperative (e.g. signaling molecules) to antagonistic (e.g. antibiotics) (16). Some exometabolites, such as antibiotics, are known to increase in production upon entry into stationary phase (6). In addition, computational models have predicted that costless exometabolite production, such as central carbon compounds, may be common among microbes (17). Untargeted exometabolomic profiling approaches have recently become popular because of advances in the sensitivity and throughput of mass spectrometers (18). This approach provides an experimental basis to observe the breadth of exometabolites produced by microbial strains and strain-specific contributions to the exometabolite pool. Characterizing the exometabolite profile of a microbe over time is a useful approach to understand the dynamic interplay between cell physiology and their environment.

We present an investigation of three environmental bacterial strains that are commonly found associated with terrestrial environments (soils or plants) (Table 1). These strains were chosen because of reported (19) and observed interactions in the lab. Our previous work established a robust and flexible approach to investigate microbial exometabolite production in either monoculture or co-culture (20). Our approach uses filter plates that allow for the separation of cells from an exometabolite reservoir. Here, we examined the detailed exometabolite and transcript dynamics for each of these three environmental strains over stationary phase, and identified key features of their exometabolomic responses to resource limitation. We found that exometabolite production is dynamic through stationary phase, and that accumulated exometabolites were likely released from intact cells. We also found that a majority of released exometabolites were strain-specific, suggesting that different bacterial strains have individualized responses to resource limitation. Finally, we found that all three strains re-routed metabolic flux in stationary phase.

**Table 1.** Bacterial strains used in study.

|  |  |  |  |
| --- | --- | --- | --- |
|  | *Burkholderia thailandensis* E264 (60) | *Chromobacterium violaceum* ATCC 31532 (61) | *Pseudomonas syringae* pathovar *tomato* DC3000 (62) |
| Family | Burkholderiaceae | Neisseriaceae | Pseudomonadaceae |
| Genome size | 6.72 Mb | 4.75 Mb | 6.53 Mb |
| ORFs | 5,641 | 4,371 | 5,853 |

## Results

### Each strain had a distinct exometabolite profile in stationary phase

In total, 10,352 features were detected by mass spectral analysis (Table 2, Supplementary Fig. S1) across the three strains. These features represent what we defined as released exometabolites (see Methods: Mass spectrometry analysis section). Polar analysis in positive mode detected the most features (5,077 features) followed by polar analysis in negative mode (2,481 features), nonpolar analysis in negative mode (1,399 features), and nonpolar analysis in positive mode (1,395 features). Most features were strain-specific, and the number of unique features from any one strain outnumbered the total number of features shared by at least two strains (1494 features, ~16.9%). Of the 1494 shared features, ~12.7% were shared among all three strains. Specifically, *B. thailandensis* had the most unique detected features (~41.8%), followed by *P. syringae* (~25.2%) and *C. violaceum* (~18.6%) compared to all detected features. These data suggest that, despite growth in minimal medium initially containing one carbon source, an abundance of strain-specific exometabolites are produced during stationary phase.

**Table 2:**
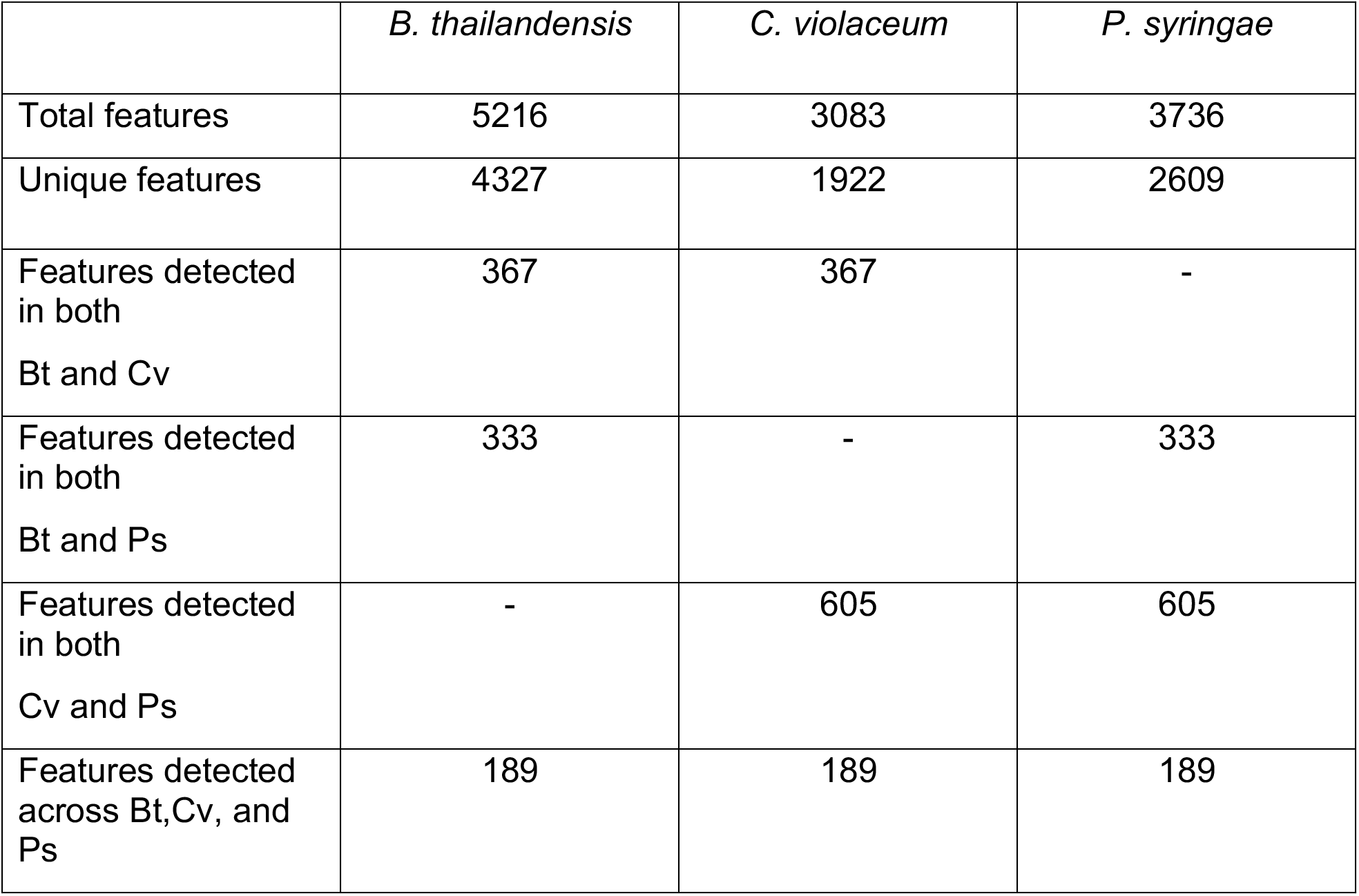
Summary of released exometabolites for each strain. Bt is *B. thailandensis*, Cv is *C. violaceum*, and Ps is *P. syringae*.

### Exometabolite composition and their temporal dynamics drive profile differences between strains

We were interested in understanding the stability of exometabolite profiles over stationary phase (Fig. 1). Each strain had strain-specific exometabolite profiles (global Adonis 0.342 ≤ *r*^2^ ≤ 0.394, *P* value ≤ 0.001, all pair-wise FDR-adjusted *P* values ≤ 0.006) and directional temporal dynamic in their exometabolites across all polarity/ionization modes. For each strain, exometabolite profiles from exponential growth phase were distinct from stationary phase profiles. Temporal trajectories in exometabolite profiles were highly reproducible for each strain across biological replicates (PROTEST; all pairwise *r*^2^ values ≥ 0.766, 0.924, and 0.701, average *P* value = 0.001, 0.006, and 0.028, and max *P* value = 0.067, 0.022, and 0.083 for *B. thailandensis, C. violaceum*, and *P. syringae*, respectively). Separately, strain and time were able to explain some variation in the dataset but, across all polarity/ionization modes, the interaction effect of strain x time explained the most variation (Supplementary Table S1). Only shared exometabolites were included in the generation of the PCoA because inclusion of all exometabolites led to strong strain-specific (variation explained ≥ 0.890) patterns (Supplementary Fig. S2). This was expected given the large number of unique features detected for each strain (Table 1). Focusing on exometabolites shared by at least two strains allowed us to discern strain-specific and temporal patterns, and still provided statistically significant differences in exometabolic profiles.

**Figure 1.**
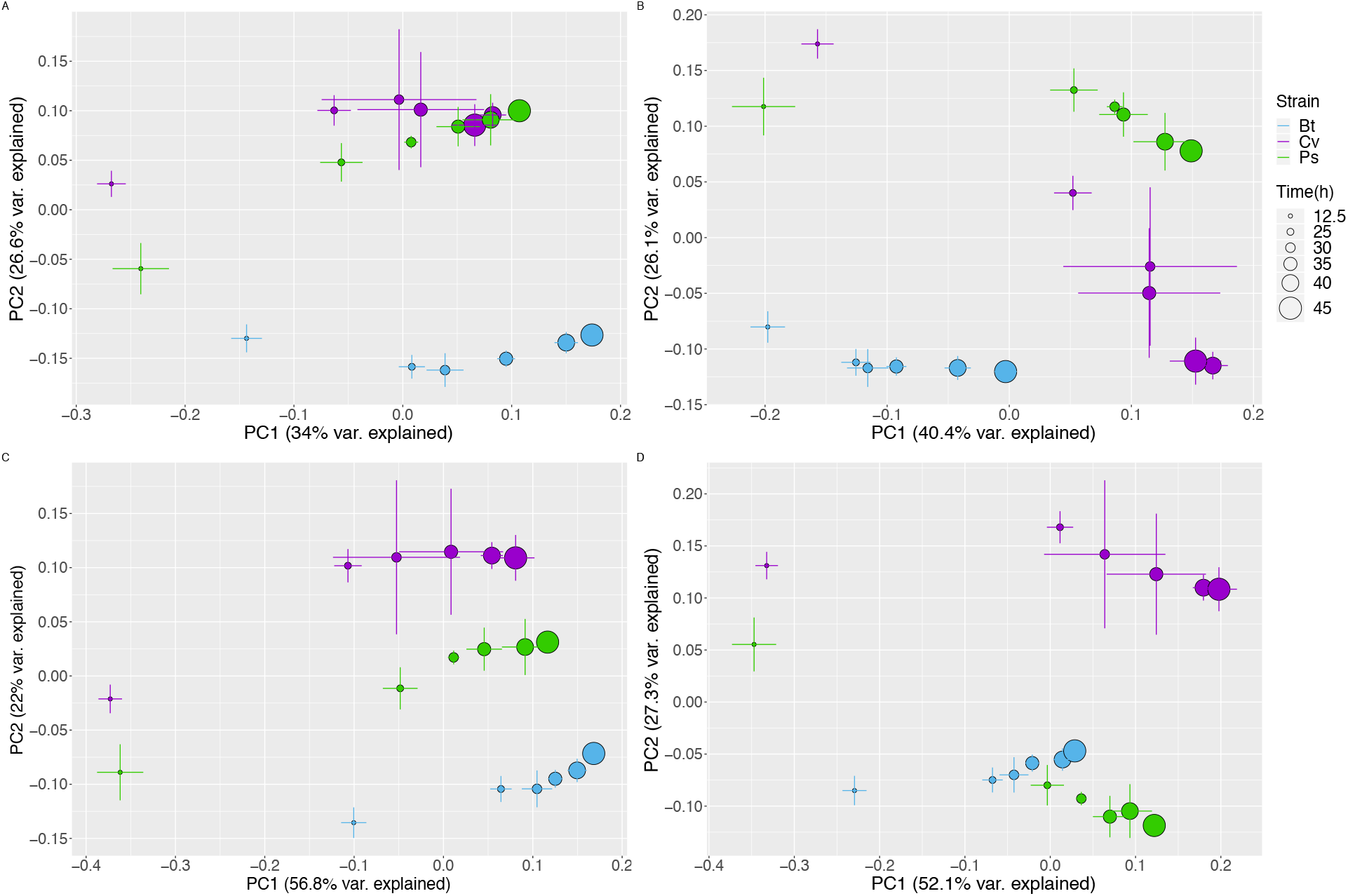
Exometabolite profiles differ by strain and time. PCoA plots for Polar positive (top left), polar negative (top right), nonpolar positive (bottom left), and nonpolar negative (bottom right) for exometabolites that were shared by at least two strains. Strain is indicated by color (*B. thailandensis* (blue), *C. violaceum* (purple), *P. syringae* (green)) and time is indicated by the size of the point. Error bars are 1 standard deviation around the mean axis scores of n = 2 to 4 replicates destructively sampled from the same time point/strain condition.

Similarly, hierarchal clustering analysis revealed strain-specific features and their dynamics (Supplementary Fig. S3). Most features reached maximum accumulation in late stationary phase. Notably, exometabolites accumulated despite generally steady strain population levels (Supplementary Fig. S4). We did observe ~1 generation in *B. thailandensis* and *P. syringae* over the course of stationary phase but this took 20 h to complete. Largely consistent population levels and a lack of death phase suggest that many exometabolites were released by intact cells rather than by lysis. To add support, transcriptomics data indicate multiple organic molecule transporters were either consistently expressed throughout the time series or differentially expressed (Supplementary Table S2, Supplementary Dataset 5). In summary, despite growth arrest, each strain continued to produce (and the media accumulates) a distinctive and dynamic profile of exometabolites well into stationary phase.

### Identity of stationary phase exometabolites

Of the total set of exometabolite features, only 188 (~1.8%) could be identified (Fig. 2, Supplementary Figs. S5 & S6). These were classified according to the Metabolomics Standards Initiative (MSI): MSI level 1 (Identified compounds) and MSI level 2 (putatively identified compounds). Most of the identified exometabolites were uniquely produced by one strain under our experimental conditions, though there were some exometabolites shared across strains, particularly between *C. violaceum* and *P. syringae* (Supplementary Fig. S7, Supplementary Dataset 4). Many of the identified exometabolites, particularly those molecules involved in central metabolism, such as amino acids, nucleotides/nucleosides, and carboxylic acids, were classified using an in-house standard in accordance with MSI level 1. In addition, MSI level 1 exometabolites such as ectoine, proline, trehalose, and glutamate likely indicated a cellular stress (e.g. osmotic stress).

**Figure 2.**
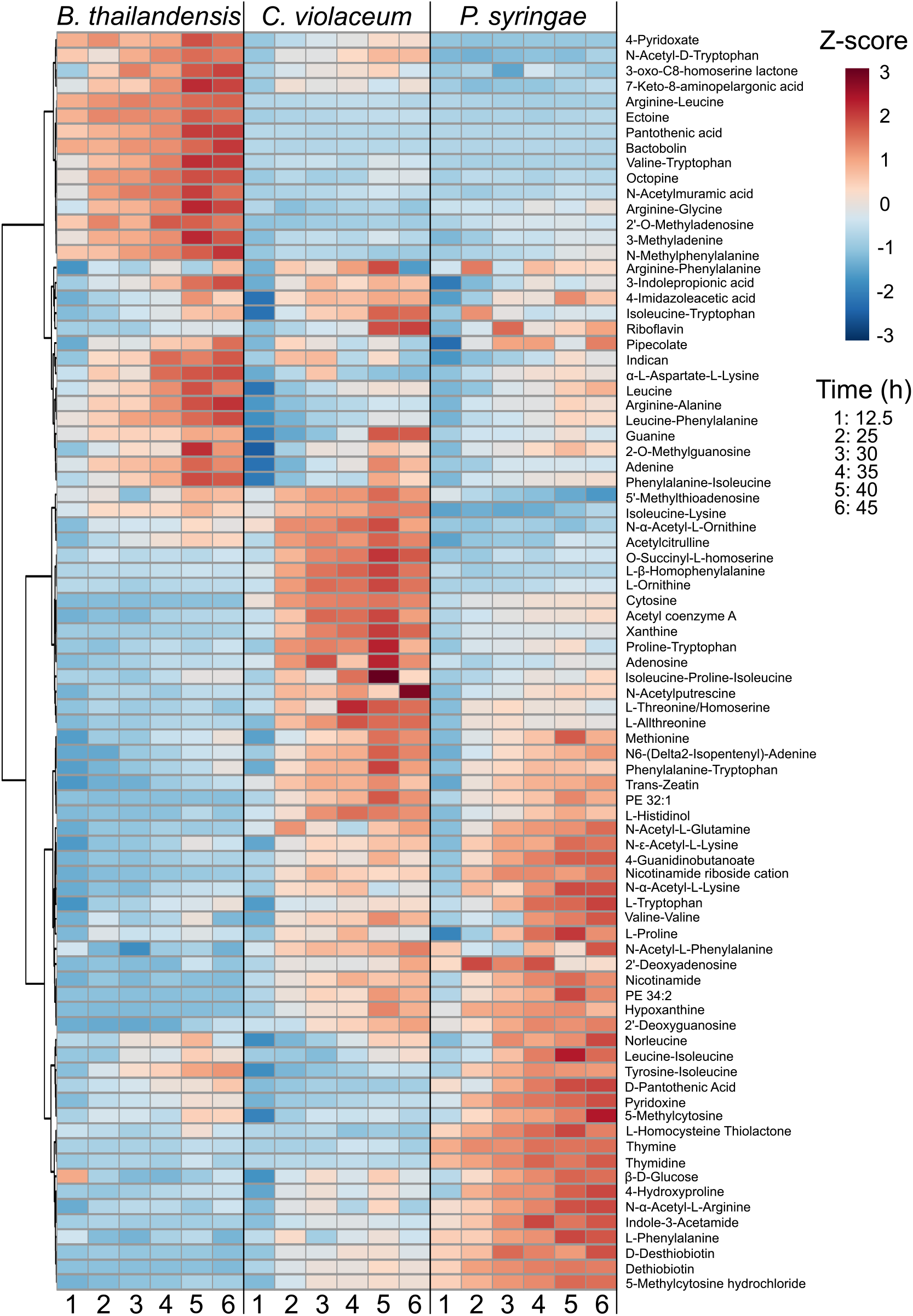
Released and identified exometabolites and their temporal dynamics. A heat map of identified exometabolites in polar positive mode is shown, where samples are columns are exometabolites are in rows. Each sample is the average of independent time point replicates (n = 3 or 4). Euclidean distance was calculated from Z-scored mass spectral profiles. Features were clustered by Ward’s method.

Exometabolites putatively identified at MSI level 2 were annotated by matching MS/MS fragmentation to a reference database. MSI level 2 exometabolties included secondary metabolites such as bactobolin, yersiniabactin, and acyl homoserine lactones (AHLs) produced by *B. thailandensis, P. syringae*, and *C. violaceum*, respectively. Bactobolin and yersiniabactin are bioactive molecules, previously characterized as a bacteriostatic antibiotic (21) and a siderophore/virulence factor (22), respectively. AHLs induce quorum sensing in *C. violaceum*, and are linked to the production of hydrogen cyanide, antibiotics, and proteases (23, 24). These putatively identified secondary exometabolites suggest that resource limitation primed strains for competition via chemical warfare or nutrient scavenging. These data also suggest that a competitive phenotype may be standard among bacteria even in the absence of non-kin competitors, suggesting either priming for interspecific competition or engagement in intraspecific competition. Finally, a large proportion of putatively identified exometabolites were dipeptides, suggesting either the degradation of proteins (25) or the formation of dipeptides.

### Annotation of remaining unidentified exometabolites

To maximize annotation of remaining unidentified MS/MS data, we performed chemical ontology analysis to determine chemical classes of exometabolites produced in stationary phase. Using *in silico* prediction of exometabolites by MS/MS fragmentation patterns, we putatively characterized compound classes (MSI level 3 designation). Broadly, carboxylic acids and derivatives were the most abundant type of exometabolite produced in stationary phase for all strains (Fig. 3A). This is expected because carboxylic acid derivatives are prominent in cellular constituents and molecules involved in primary metabolism (e.g. TCA cycle). However, MSI level 3 exometabolites revealed considerable quantification of molecules related to fatty acyls, organonitrogen compounds, organooxygen compounds, and benzene and substituted derivatives, suggesting additional classes of exometabolites contributing to the exometabolite pool that are unable to be identified by MSI level 1 and level 2 standards. These chemical ontologies were resolved further to the direct parent level (Fig. 3B). Amino acids and peptides were the most abundant and common exometabolites across all identification levels. In particular, dipeptides were the most abundant exometabolite. Transcriptomics data also indicated that dipeptide transporters for each strain were either consistently expressed or differentially expressed over time (Supplementary Dataset 5, Supplementary Table S2).

**Figure 3.**
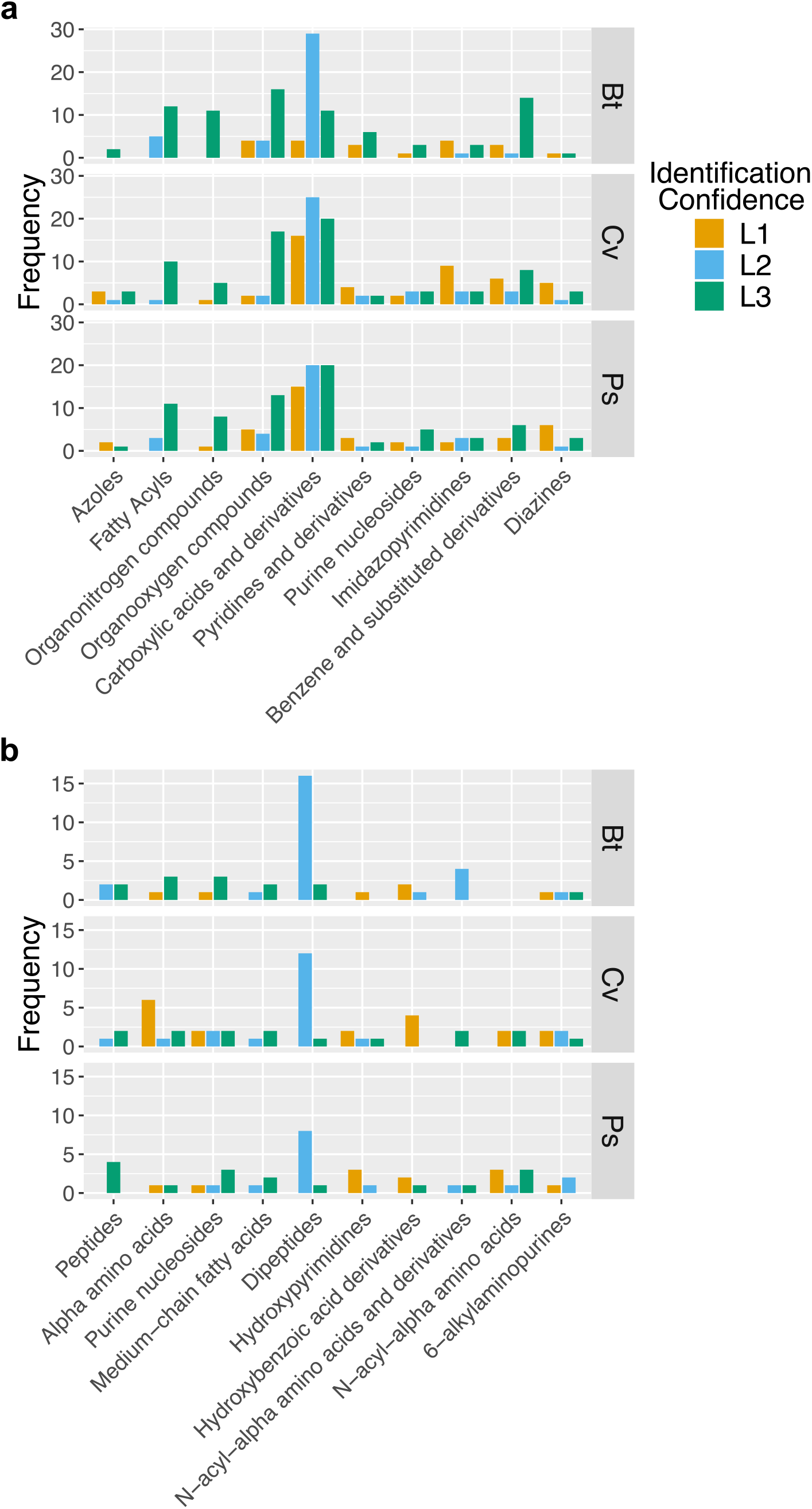
Chemical ontologies at different MSI levels. ClassyFire was used to categorize identified (MSI level 1 and level 2) and *in silico* predicted MS/MS data (MSI level 3) at the a) class and b) direct parent levels. The top ten chemical ontologies are provided for each classification level.

### Insights into stationary phase metabolic re-routing

We then aimed to interpret strain metabolism in stationary phase by focusing on exometabolites most confidently identified (MSI level 1). Pathway analysis identified similarities in the enrichment of cellular pathways across strains (Supplementary Dataset 6). For example, all strains were enriched in purine metabolism, both *B. thailandensis* and *P. syringae* were enriched in vitamin B6 metabolism, and, both *C. violaceum* and *P. syringae* were enriched in pyrimidine metabolism, tryptophan metabolism, sulfur metabolism, lysine biosynthesis, arginine biosynthesis, arginine and proline metabolism, and aminoacyl-tRNA biosynthesis. Pathways enriched that were unique to a strain included amino acid metabolisms, butanoate metabolism, glyoxylate and dicarboxylate metabolism, glutathione metabolism, and pyruvate metabolism in *P. syringae* and, cysteine and methionine metabolism in *C. violaceum*. Taken together, temporal dynamics in exometabolites led to the enrichment of various metabolic pathways, with the most prominent alterations occurring in *P. syringae*.

For each strain, we examined the 10 most abundant exometabolites that accumulated and detected at last time point (45 h) for positive and negative polar exometabolites. We included all MSI level 1 exometabolites in this analysis. A majority of the most abundant exometabolites were unique to each strain (Fig. 4), and strain-specific exometabolites were more abundant in comparison to the other strains (ANOVA FDR-adjusted *P* values ≤ 0.01) with the exception of 5’-methylthioadenosine and hypoxanthine. The majority of strain-specific abundant exometabolites suggest that each strain differs in the regulation of major metabolic pathways. This differential regulation can reflect direct influence on the pathways involved in the production of an exometabolite or it can reflect influence on the transport of an exometabolite.

**Figure 4.**
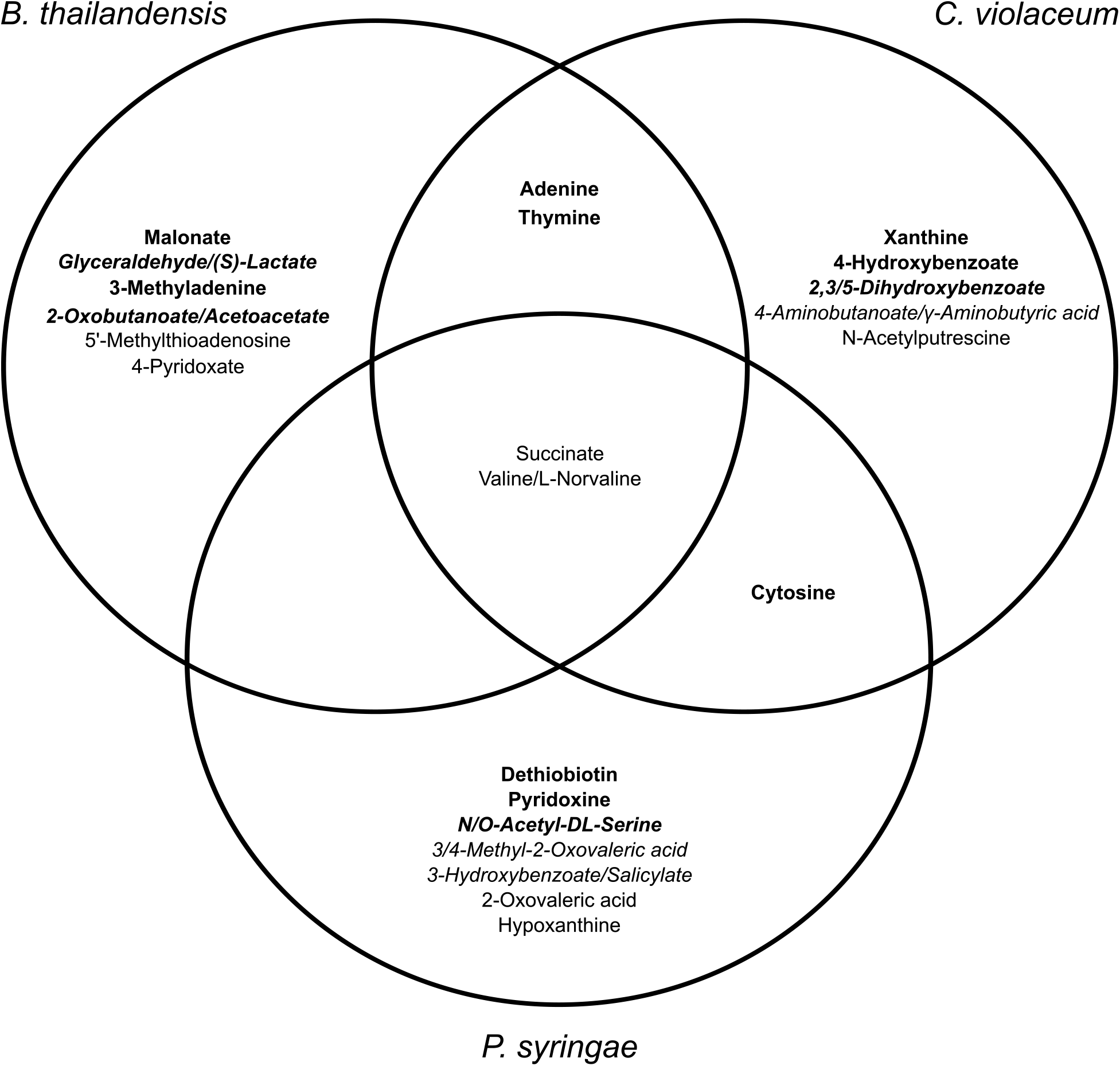
Distinctions and overlaps between the most abundant exometabolites in each strain. Exometabolites in bold passed criteria for “released”. Exometabolites in italics are isomers and could not be resolved to determine the exact identification.

Of the most accumulated exometabolites, succinate was a common exometabolite detected in all strains, and this is unsurprising as it is directly involved in central metabolism. We overlaid temporal log fold changes in gene expression onto KEGG pathways involved in succinate production (Fig. 5). These data suggest that all strains re-routed metabolism during stationary phase. For the most part, enzymes involved in glycolysis and the TCA were downregulated in all strains (Supplementary Figs. S8 & S9). With regard to succinate production, both *B. thailandensis* and *C. violaceum* appear to have re-routed metabolism to use the glyoxylate cycle, as supported by the upregulation of isocitrate lyase and upregulation of enzymes involved in the ß-oxidation of fatty acids. Quorum-sensing mediated metabolic re-routing to the glyoxylate cycle during stationary phase in *B. thailandensis and Burkholderia glumae* was found to combat alkalinity toxicity (26, 27). Furthermore, the greatest upregulation in isocitrate lyase was observed in *Burkholderia cenocepacia* during stationary phase compared to other abiotic stressors (28). This supports the notion that a re-routing metabolism to the glyoxylate cycle in stationary phase may be a shared feature among members of the genus *Burkholderia. P. syringae* appears to have re-routed metabolism to use the methylcitrate cycle to generate succinate, as evidenced by the upregulation of 2-methylisocitrate lyase. Other sources of succinate production include the GABA shunt (*B. thailandensis* and *P. syringae)*, succinyl-CoA:acetate CoA-transferase (*B. thailandensis* and *P. syringae*), and fumurate reductase (*C. violaceum*). In all strains, stationary phase results in dynamic exometabolite and transcriptional changes that supported maintenance of cell integrity under resource-limited conditions.

**Figure 5.**
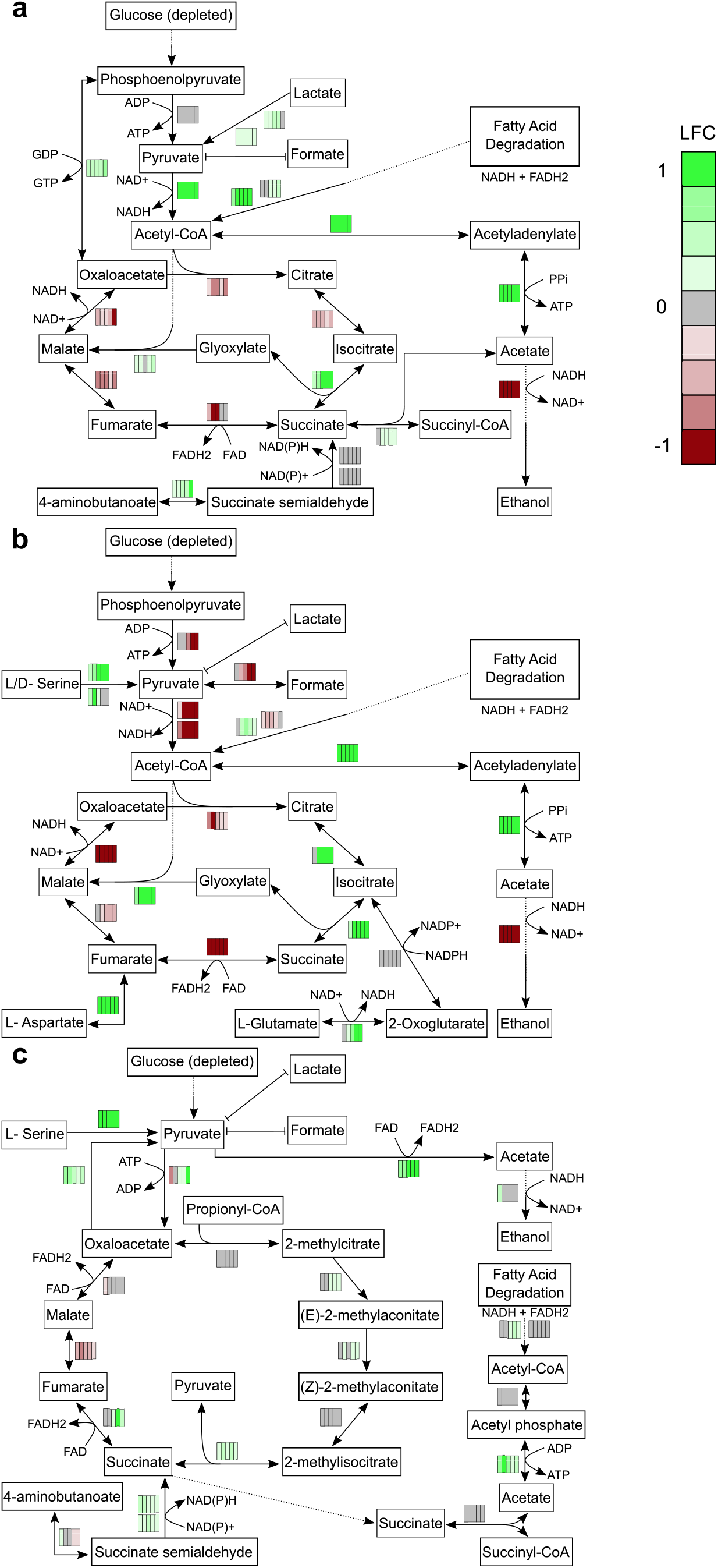
Temporal changes in transcriptomics reveal re-routing of metabolic flux towards succinate production. Log_2_-fold change (LFC) values were mapped onto pathways involved in succinate production for a) *B. thailandensis*, b) *C. violaceum*, and c) *P. syringae*. LFC values are represented by rectangles alongside each reaction in the pathway map. Each column represents the 5 stationary phase time points. Colors within each rectangle represent LFC (green-upregulated, red-downregulated) compared to the exponential phase time point.

## Discussion

Most microbes in the environment (e.g. soil, open ocean) exists predominantly in stationary phase or in dormancy. Non-growth states can result from resource limitation in a local environment with inaccessible or depleted nutrients. We studied exometabolite production in stationary phase across three separate bacterial strains. We have found that resource limitation results in dynamic and directional exometabolite profiles even after the cells entered stationary phase. Exometabolite dynamics were observed across all three strains (Fig. 1 & Supplementary Fig. S3). We specifically focused our analyses on released exometabolites. That is, exometabolites that accumulated in the medium over time. Even though we applied a very conservative definition to identify features that accumulated, we detected and characterized thousands of features that met our criteria. However, in the end, only a subset of these features could be identified using standards, MS/MS databases, and computational predictions based on chemical characteristics (Fig. 2 & Supplementary Fig. S5 & S6, Supplementary Dataset 4).

Exometabolites could accumulate over stationary phase by two mechanisms. First, exometabolites could be transported passively or actively across the cell membrane. Second, cells could lyse and spill primary metabolites and other debris into the media (29). Our results suggest that a major contributing factor to exometabolite accumulation for all three strains investigated here was exometabolite release from intact cells. In fact, we did not observe a death phase over stationary phase (Supplementary Fig. S4). Live cells generally remained at consistent levels throughout stationary phase. One generation during stationary phase was observed for both *B. thailandensis* and *P. syringae.* Given the downregulation of multiple enzymes in central metabolism (Supplementary Figs. S8 & S9), this generation was likely the result of reductive cell division (30). Additional evidence of exometabolite release from intact cells was provided by RNAseq analysis. Transcriptomics results indicate the upregulation or consistent expression of transporters (Supplementary Dataset 5). In a previous study, Paczia *et al.* also observed similar patterns of exometabolite accumulation in stationary phase in various strains. They were able to rule out lysis and determine that passive or active diffusion could explain exometabolite production in growth limited conditions (31). Findings from our study and from Paczia *et al.* are further supported by metabolic models that found the extracellular accumulation of central metabolites may be attributable to costless metabolic secretions in resource poor environments (17). Unintuitively, exometabolite production, particularly central carbon intermediates, by viable cells may be a common adaptation of all microorganisms under resource limitation.

In addition to the characterization of exometabolites implicated in cooperative interactions (e.g. central carbon intermediates or AHLs), we also identified exometabolites implicated in competition. An antibiotic (bactobolin) was produced by *B. thailandensis* and a siderophore (yersiniabactin) was produced by *P. syringae*, representing interference (direct harm to neighbors) and exploitative (indirect negative interaction) competition strategies, respectively (32, 33). These exometabolites are involved in interspecies competition but were still produced despite growth in monoculture. While we did not identify an exometabolite in *C. violaceum* involved in competition, we did identify AHLs, which are linked to the production of competitive exometabolites (23, 24). Taken together, stationary phase altered metabolic regulation that lead to the production of both cooperative and competitive exometabolites. Simultaneous production of both cooperative and competitive exometabolites may be an advantageous strategy to sustain kin while maintaining competition for scarce resources. Additional studies are needed to understand the interplay between cooperation and competition under changing environmental conditions.

Putative (MSI level 2) exometabolite identifications provided evidence for the release of dipeptides (Fig. 3B) and transcriptomics provided evidence for differentially regulated or consistent expression of dipeptide transporters (Supplementary Dataset 5). Hydrolysis by dipeptidyl peptidases of ribosomal proteins or degradation of other polypeptide chains can be one source of dipeptide production. Estimates in *E. coli* have shown that 50-80% of ribosomes were degraded upon transition from exponential phase to stationary phase (34). Interestingly, another source of dipeptides may be attributed to active production. Recent studies have examined dipeptide formation by adenylation domains in nonribosomal peptide synthetases (NRPS) (35, 36). All strains in our study have numerous NRPS that could contribute to the production of dipeptides (data not shown). Furthermore, one dipeptide was characterized as a cyclic dipeptide. Cyclic dipeptides can be involved in cell communication (37). Thus, the versatility of chemical ecology facilitated by dipeptides makes it important to understand how dipeptides are formed and characterize the environments that induce their production.

Efforts have been put forth to annotate all MS/MS data (38). We used the same approach to computationally predict and classify the chemical ontology of MS/MS data not identified at MSI level 1 or level 2 (Fig. 3). Differences between *in silico* predictions of MS/MS data (MSI level 3) and MSI levels 1 and 2 was most apparent at the class level (Fig. 3A). This knowledge can be used to direct research efforts and analytical techniques to identify underrepresented classes of compounds in experimental systems.

Pathway analysis revealed overlaps across all strains in the enrichment of nucleobase metabolic processes. This finding is consistent with a previous study in *E. coli* that observed the extracellular accumulation of nucleobases upon entry into stationary phase (11). Ribosome degradation is initiated in growth-limiting environments and is a likely source of nucleobase accumulation due to the degradation of rRNA (34). A comprehensive study of both exometabolites and intracellular metabolites levels could link the dynamic relationship between microbial metabolism and survival during nutrient stress.

Microbes in growth-arrested states can re-route metabolic flux to maintain the proton motive force (PMF) and stabilize ATP levels (8). We used a combination of exometabolomics and transcriptomics to shed light on metabolic re-routing in each strain investigated. Notably, all three strains accumulated high levels of succinate, and this was further supported by transcript data that showed upregulation of pathways involved in succinate production (Fig. 5). We found that the major metabolic re-routing in stationary phase included transitioning to the glyoxylate cycle in *B. thailandensis* and *C. violaceum* and to the methylcitrate cycle in *P. syringae.* While we were not able to characterize transporters, we did find that C4-dicarboxylic acids were transcriptionally active in all three strains (Supplementary Dataset 5). It could be that succinate export is facilitated by a succinate/proton symporter for maintenance of the PMF. However, both cycles involved in succinate production do not generate ATP. Stable ATP levels are also necessary to maintain cell viability. Transcriptomic evidence from all three strains suggest that ATP could be generated through the production acetate. Though, this does not exclude the possibility that acetate is consumed for the generation of acetyl-CoA. Combining exometabolomic and transcriptomic approaches provided increased biological interpretation that could not have been achieved by either approach in isolation.

Increased studies on growth-limited states will continue to shed light on this predominant microbial physiological state. Even in a simple minimal medium, each strain we studied dynamically altered its environment through the production of a breadth of exometabolites. A limit to our study is the lack of annotated exometabolites. Continued advances in mass spectrometry will help to annotate unknown exometabolites to reveal the full extent of exometabolites produced by a microbe. In addition, multi-omic approaches can increase biological insights and inform metabolic models. Combinations of experimental and computational approaches will be useful strategies to manage and predict the response of microorganisms to stress.

## Materials and Methods

### Bacterial strains and culture conditions

Glycerol stocks of *B. thailandensis, C. violaceum*, and *P. syringae* (Table 1) were plated on half-concentration Trypticase soy agar (TSA50) at 27°C for at least 24 h. Strains were inoculated in 7 ml of M9-0.2% glucose medium and grown for 16 h at 27°C, 200 rpm. Cultures were then back-diluted into 50 ml M9–0.2% glucose medium such that exponential growth phase was achieved after 10 h of incubation at 27°C, 200 rpm. Strains were back-diluted in 50 ml M9–0.067% glucose medium to target ODs (*B. thailandensis* 0.3 OD, *C. violaceum:* 0.035 OD, *P. syringae* 0.035 OD) such that stationary phase growth after approximately 24 h of incubation in filter plates.

### Filter plate experiments

We used the filter plate system to study each strain in monoculture over the course of stationary phase. Filter plate preparation was performed as previously described (20). Briefly, we used sterile filter plates with 0.22-μm-pore polyvinylidene difluoride (PVDF) filter bottoms (MultiScreen GV Filter Plate, 0.22 μm, MSGVS2210, Millipore). Prior to use, filter plates were washed three times with sterile water using a vacuum apparatus (NucleoVac 96 vacuum manifold; Clontech Laboratories). The filter of well H12 was removed with a sterile pipette tip and forceps, and 31 ml of M9–0.067% glucose medium was added to the reservoir through well H12. Each well was then filled with 130 μl of back-diluted culture in M9–0.067% glucose medium or medium only. For each plate, a custom R script (RandomArray.R [see the GitHub repository]) was used to randomize the placement of strains in the wells so that each strain occupied a total of 31 wells per plate. Each monoculture time course was independently replicated four times for a total of 12 experiments. The time course included 6 time points: an exponential phase point (12.5 h) and 5 points assessed every 5 h over stationary phase (25 h – 45 h). Plates were destructively sampled, comprising a total of 72 plates for the entire experimental design of 3 strains x 6 timepoints x 4 replicates.

Filter plates were incubated at 27°C with gentle shaking (~0.32 rcf). We again used our RandomArray.R script to randomize wells used for RNA extraction (16 wells, pooled per plate) and flow cytometry (5 wells, pooled per plate). During destructive sampling, first, the wells containing spent culture assigned to RNAseq were pooled into a 1.5 mL microcentrifuge tube, flash frozen in liquid nitrogen, and stored at −80°C for RNA extraction. Next, wells containing spent culture assigned to flow cytometry were pooled, and then 20 μL was initially diluted in 180 μL Tris-buffered saline (TBS; 20 mM Tris, 0.8% NaCl [pH 7.4]), and then, after checking concentrations needed for accurate flow cytometry counts, diluted further in TBS to reach final dilutions of 1,300-fold, 1,540fold, and 900-fold for *B. thailandensis, C. violaceum, P. syringae*, respectively. Finally, spent medium (~31 ml) from the shared reservoir was transferred into 50 mL conical tubes, flash-frozen in liquid nitrogen and stored at −80 °C for subsequent exometabolite extraction.

### Flow cytometry

Diluted cultures were stained with the Thermo Scientific LIVE/DEAD BacLight bacterial viability kit at final concentrations of 1.5 μM Syto9 (live stain) and 2.5 μM propidium iodide (dead stain). Two hundred microliters of stained cultures were transferred to a 96-well microtiter U-bottom microplate (Thermo Scientific). Twenty microliters were analyzed on a BD Accuri C6 flow cytometer (BD Biosciences) at a fluidics rate of 66 μl/min and a threshold of 500 on an FL2 gate. The instrument contained the following optical filters: FL1-533, 30 nm; FL2-585, 40 nm; and FL3, 670-nm longpass. Data were analyzed using BD Accuri C6 software version 1.0.264.21 (BD Biosciences).

### Metabolomics

#### Mass spectrometry analysis

Both MS and MS/MS data were used for untargeted metabolomics analysis. A total of 257/288 metabolomic samples were used for analysis; 30 samples were removed due to failed injection and 1 sample was removed due to low intragroup reproducibility in polar analysis (Pearson’s r ≤ 0.14). MZmine (version 2.42) (39) was used for peak picking, aligning features across samples, and peak integration for both nonpolar and polar analyses and in both negative and positive ion mode. MZmine parameters for all analyses can be viewed in the XML files used by MZmine (Supplementary Dataset 1). For MS data, a feature by sample matrix was exported for additional feature filtering steps. For MS/MS data, the GNPS feature was used to export data in addition to performing a local spectra database search within MZmine (see Compound identification section, below).

We used filter featuring steps to identify exometabolites released from each strain in stationary phase. The feature filtering steps were performed as follows on a per-strain basis: 1) Features were removed if the max peak area was found in one of the replicates for the external control sample. 2) A noise filter: the minimum peak area of a feature from a replicate at the last time point (45 hr) needed to be 3X the maximum peak area of the same feature in one of the external control replicates. 3) Coefficient of variation (CV) values for each feature calculated between replicates at each time point needed to be less than 20% across the time series. 4) The minimum value of the average peak area needed to be observed in the first, exponential phase time point (12.5 h). 5) The log2 fold change of the average peak areas observed between the last (45 h) and first (12.5 h) timepoints needed to be greater than 1. 6) The time series abundance of a feature needed to have a Pearson correlation greater than or equal to 0.7.

Four final feature datasets from polar and nonpolar analysis in both ionization modes were analyzed in MetaboAnalyst 4.0 (40). Features were normalized by an internal standard (ITSD) reference feature (Supplementary Dataset 2) and cube root transformed. Reference features for polar analysis in positive (13C-15N-proline) and negative (13C-15N-alanine) was determined by the ITSD with the lowest CV value across all samples. The reference feature for nonpolar datasets was the ITSD 2-Amino-3-bromo-5-methylbenzoic acid (ABMBA). Heatmaps were generated in MetaboAnalyst using Ward’s clustering algorithm with Euclidean distances from Z-scored data. Normalized and transformed datasets were exported from MetaboAnalyst to generate principal coordinate analysis (PCoA) plots in R. Abundances for exometabolites that did not pass release criteria in each strain were replaced with NAs prior to distance matrix computation.

#### Compound identification

A three step process was used to identify compounds or characterize chemical ontologies(38). Identification confidence was assigned according to the Metabolomics Standards Initiative (MSI) (41). First, compounds were identified by an in-house reference library at the Joint Genome Institute (JGI). This reference library was curated to identify compounds based on m/z, retention time, and MS/MS spectra of standards. A compound passing the first two criteria were denoted MSI level 1. A compound passing all three criteria exceeded MSI level 1. All compounds at or exceeding MSI level 1 were identified using the reference library. This reference library was only available for polar analysis. Ranges for m/z and retention time values for compounds in the reference library were used to identify exometabolites from the MZmine analysis (Supplementary Dataset 3).

We made an effort to identify as many of the remaining compounds from both polar and nonpolar analyses that had MS/MS data. MS/MS data acquired during mass spec analysis were used to putatively identify compounds that matched to fragmentation patterns from libraries outside of JGI; these were assigned MSI level 2. First, MS/MS data was exported to GNPS format and analyzed in GNPS (42) to match fragmentation patterns against the NIST17 commercial library. Second, a local spectra database search was performed within MZmine using the entire compound library from MassBank of North American (MoNA- https://mona.fiehnlab.ucdavis.edu). For both approaches, compounds were putatively identified if cosine scores were 0.7 or above. A subset of the final feature datasets was created from compounds identified at MSI level 1 and level 2 (Supplementary Dataset 4). These datasets were processed in MetaboAnalyst (see Mass spectrometry analysis section, above) to generate heat maps, perform pathway analysis (see Pathway analysis section, below), and perform ANOVA analysis between strains exometabolite abundances.

All remaining unidentified compounds with MS/MS data were analyzed with CSI:Finger ID and assigned MSI level 3. This method provides the putative chemical ontology of a compound. The top CSI:Finger ID match was used for each compound. Then, lnChl keys from all MSI levels were used to perform a chemical ontology analysis using ClassyFire version 1.0. SDF files from ClassfyFire were exported from each analysis to extract both Class level and Direct Parent level ontologies. These data were then exported to R for data visualization.

#### Pathway analysis

MSI level 1 exometabolites were used for pathway analysis. First, MSI level 1 from both polar positive and polar negative modes were separately uploaded to MetaboAnalyst for normalization by a reference standard and cubed root transformed. Normalized and transformed datasets from both polarities were then combined. Isomers were then removed since their identification could not be resolved. The dataset still contained exometabolites identified in both ionization modes. To resolve this, we split the dataset into two. One dataset contained exometabolites identified in only one ionization mode and exometabolites from positive ionization mode from exometabolites that were identified in both ionization modes. The second dataset also contained exometabolites identified in only one ionization mode and, exometabolites from negative ionization mode from exometabolites that were identified in both ionization modes. Both datasets were then split into three separate datasets by strain and used to perform pathway analysis in MetaboAnalyst. The normalized and transformed concentration tables for each strain was analyzed with time set as the continuous group label. Exometabolites were mapped onto *Escherichia coli* K-12 MG1655 KEGG pathways (version Oct2019) and the global test pathway enrichment analysis algorithm was used to generate FDR-adjusted *P* values. Pathways were considered enriched if at least 3 exometabolites mapped to the pathway and, FDR-adjusted *P* values ≤ 0.05 for a given pathway in a strain across both ionization mode analyses.

### Transcriptomics

#### RNA quality filtering and differential gene expression (DGE) analysis

Count matrices for each strain were quality filtered in two steps prior to DGE: genes containing 0 counts in all samples were removed and genes with a count ≤ 10 in more than 90% of samples were removed. DGE was performed in DESeq2 version 1.22.1 (43). We tested for differential gene expression by evaluating genes that changed at any time point (FDR < 0.01). Genes with differential expression were then evaluated for log2 fold changes >1. Specifically, we focused on genes involved in transport (see Transporter analysis section, below).

#### Defining expression minimums

A cumulative abundance plot was generated for each strain by organizing locus IDs from low transcript counts to high transcript counts and plotting the % of total transcripts against the % of total read counts (44, 45). The 25^th^ quantile was calculated to obtain the transcript count value that defined a low expression minimum. That is, all genes with transcript counts above this minimum were considered to be expressed in the cell, regardless of longitudinal differential expression.

#### Transporter analysis

TransportDB 2.0 (http://www.membranetransport.org/transportDB2/index.html) was used to annotate transporters in each strain (46). Annotated transporters were then evaluated to determine differential expression or expression above the low expression minimum.

#### KEGG *pathway analysis*

We extracted log2 fold change (LFC) values from transcripts in each strain from DESeq analysis. Log2 fold change were obtained by comparing each stationary phase time point to the exponential time point 1 (12.5 h). We then mapped longitudinal LFCs onto KEGG pathways for each strain using the pathview package in R. First, K numbers were assigned to genes for both *C. violaceum* and *P. syringae* using BlastKOALA (version 2.2). K numbers were not assigned to *B. thailandensis* because KEGG identifiers were available. KEGG identifiers for *B. thailandensis* and K numbers assigned to *C. violaceum* and *P. syringae* were used to map longitudinal LFCs onto KEGG pathways. Pathways of interest were curated and manually edited in Inkscape (verision 0.92.4) using a colorblind palette.

## Code availability

Computing code and workflows are available at https://github.com/ShadeLab/Paper_Chodkowski_MonocultureExometabolites_2020. R packages used during computing analyses included vegan (47), ggplot2 (48), VennDiagram (49), pairwiseAdonis (50), patchwork (51), DESeq2 (43), pathview (52), KEGGREST (53), and helper functions (54–57).

## Data availability

Genomes for *B. thailandensis, C. violaceum*, and *P. syringae* are available at JGI Genome Portal under accession numbers 637000052, 2724679652, and 2508501074, respectively JGI ProposaI ID 502921. An improved annotated draft genome of *C. violaceum* is available under NCBI Project ID 402426 (Genbank Accession ID: PKBZ00000000). Re-sequencing efforts for *B. thailandensis* and *P. syringae* are under NCBI project IDs 402425 and 402424, respectively. Metabolomics data and transcriptomics data are also available at JGI Genome Portal (59) under JGI Proposal ID 502921. Large data files (e.g. MZmine project files) are available upon request.

## Author contributions statement

J.C. and A.S. conceived of and designed the study. J.C. performed the research and analyses. J.C. and A.S. wrote the manuscript.

## Acknowledgments

This material is based upon work supported by the National Science Foundation under Grant No DEB#1749544, by Michigan State University, and by a DOE-JGI Community Science Program award (Proposal ID 502921). The work conducted by the U.S. Department of Energy Joint Genome Institute, a DOE Office of Science User Facility, is supported under Contract No. DE-AC02-05CH11231. J.C. was supported by the Eleanor L. Gilmore Fellowship from the Department of Microbiology and Molecular Genetics. We thank Katherine B. Louie and Benjamin P. Bowen for support in mass spectral analysis.

## Additional information

Competing interest statements

The authors declare no competing interests

## Supplemental files

**Supplementary Dataset 1**. XML files containing MZmine parameters for running analysis in batch. This includes three files (MS analysis, MSMS analysis, and MSMS spectral match) for each nonpolar negative, nonpolar positive, polar negative, and polar positive modes.

**Supplementary Dataset 2**. Mass spectrometry metadata for internal standards from polar analysis.

**Supplementary Dataset 3**. Mass spectrometry metadata of JGI in-house standards from polar analysis.

**Supplementary Dataset 4**. MSI level 1 and level 2 exometabolites that fit criteria for released in each strain.

**Supplementary Dataset 5**. Transcriptomics analysis of transporters in each strain.

**Supplementary Dataset 6**. Exometabolite pathway analysis results.

**Supplementary Dataset 7**. Metadata for each sample injection onto the mass spectrometer.

